# Antibiotic interactions shape short-term evolution of resistance in *E. faecalis*

**DOI:** 10.1101/641217

**Authors:** Ziah Dean, Jeff Maltas, Kevin B. Wood

## Abstract

Antibiotic combinations are increasingly used to combat bacterial infections. Multidrug therapies are a particularly important treatment option for *E. faecalis*, an opportunistic pathogen that contributes to high-inoculum infections such as infective endocarditis. While numerous synergistic drug combinations for *E. faecalis* have been identified, much less is known about how different combinations impact the rate of resistance evolution. In this work, we use high-throughput laboratory evolution experiments to quantify adaptation in growth rate and drug resistance of *E. faecalis* exposed to drug combinations exhibiting different classes of interactions, ranging from synergistic to suppressive. We identify a wide range of evolutionary behavior, including both increased and decreased rates of growth adaptation, depending on the specific interplay between drug interaction and drug resistance profiles. For example, selection in a dual β-lactam combination leads to accelerated growth adaptation compared to selection with the individual drugs, even though the resulting resistance profiles are nearly identical. On the other hand, populations evolved in an aminoglycoside and β-lactam combination exhibit decreased growth adaptation and resistant profiles that depend on the specific drug concentrations. We show that the main qualitative features of these evolutionary trajectories can be explained by simple rescaling arguments that correspond to geometric transformations of the two-drug growth response surfaces measured in ancestral cells. The analysis also reveals multiple examples where resistance profiles selected by drug combinations correspond to (nearly) optimized linear combinations of those selected by the component drugs. Our results highlight trade-offs between drug interactions and resistance profiles during the evolution of multi-drug resistance and emphasize evolutionary benefits and disadvantages of particular drug pairs targeting enterococci.

## INTRODUCTION

The rapid rise of antibiotic resistance poses a growing threat to public health (1, 2). The discovery of new antimicrobial agents is a long and difficult process, underscoring the need for new approaches that optimize the use of currently available drugs. In recent years, significant efforts have been devoted to designing evolutionarily sound strategies that balance short-term drug efficacy with the long-term potential to develop resistance. These approaches describe a number of different factors that could modulate resistance evolution, including interactions between bacterial cells (3, 4, 5, 6, 7, 8), synergy with the immune system (9), spatial heterogeneity (10, 11, 12, 13, 14, 15), epistasis between resistance mutations (16, 17), precise temporal scheduling (18, 19, 20, 21), and statistical correlations between resistance profiles for different drugs (22, 23, 24, 25, 26, 27, 28, 29, 30, 31).

Drug combinations are an especially promising and widely used strategy for slow-ing resistance (32). From a clinical perspective, synergistic interactions–where the combined effect of the drugs is greater than expected based on the effects of the drugs alone (33)–have long been considered desirable because they provide strong antimicrobial effects at reduced concentrations. By contrast, drug pairs that interact antagonistically–effectively weakening one another in combination–have been traditionally avoided. Work over the last decade has challenged this conventional wisdom by demonstrating that synergistic interactions have a potentially serious drawback: they may accelerate the evolution of resistance (34, 35, 36). Similarly, antagonistic interactions can slow or even reverse the evolution of resistance (37). These results indicate that drug interactions underlie a natural trade-off between short-term efficacy and long-term evolutionary potential (38). In addition, recent work has shown that cross-resistance (or collateral sensitivity) between drugs in a combination may also significantly modulate resistance evolution (27, 39, 26). As a whole, these studies show that drug interactions and collateral effects may combine in complex ways to influence evolution of resistance in multi-drug environments.

Combination therapies are an important tool for treating enterococcal infections, which lead to significant morbidity and mortality (40, 41, 42, 43). *E. faecalis* is among the most commonly isolated enterococcal species and underlies a host of human infections, including infective endocarditis, infections of the urinary tract or blood stream, and infections related to surgical devices and medical implants. Multiple combination therapies have been proposed or are currently in use for *E. faecalis* infections (44, 45, 40). While synergistic combinations are the standard–particularly for high inoculum infections–relatively little is known about how different combinations affect the potential for, and rate of, resistance evolution.

To address these questions, we use large scale laboratory evolution to measure growth adaptation and phenotypic resistance in populations of *E. faecalis* exposed to four different two-drug combinations over multiple days. The drug pairs include several clinically relevant combinations–for example, two β-lactams or a β-lactam and an amimoglycoside–and exhibit a range of interactions, from synergistic to strongly antagonistic (suppressive). Our results reveal rich and at times surprising evolutionary behavior. In all cases, we find that different dosing combinations lead to significantly different rates of growth adaptation, even when the level of inhibition is constant. In some cases, differences in growth adaptation appear to be driven by selection for distinct cross-resistance profiles, while in other cases, strong interactions between the drugs lead to different adaption rates but highly similar profiles. We show that qualitative features of these evolutionary trajectories can be understood using simple rescaling arguments that link resistance profiles in evolving populations to geometric transformations of the two-drug response surface in ancestral cells. Our results represent a quantitative case study in the evolution of multidrug resistance in an opportunistic pathogen and highlight both potential limitations and unappreciated evolutionary benefits of different drug combinations.

## RESULTS

### Selection of antibiotic pairs with different interactions

We first set out to identify a set of two-drug combinations that include a range of interaction types: synergistic, antagonistic, and suppressive (i.e. strongly antagonistic so that the effect of the combination is less than that of one of the drugs alone). To do so, we measured the per capita growth rate of *E. faecalis* V583 populations in liquid cultures exposed to multiple drug pairs at 90-100 dosage combinations per pair. We estimated per capita growth rate in early exponential phase from optical density (OD) time series acquired using an automated microplate reader and plate stacker (Methods). The type of interaction for each drug pair is defined by the shape of the contours of constant growth (“isoboles”) describing the growth response surface *g* (*D*_1_, *D*_2_), where *D*_*i*_ is the concentration of drug *i*. Linear contours of constant growth represent additive (non-interacting) pairs–for example, the effect of one unit of drug 1 or drug 2 alone is the same as that of a combination with half a unit of each drug. Deviations from additivity include synergy (antagonism), which corresponds to contours with increased concavity (convexity), rendering an equal mixture of the two drugs more (less) effective than in the non-interacting scenario. While other metrics exist for quantifying drug interactions (see, for example, (33)), we choose this Loewe null model (46) because, as we will see, its simple geometric interpretation provides useful intuition for interpreting evolutionary trajectories (37, 35). Based on these interaction measures and with an eye towards choosing clinically relevant combinations when possible, we decided to focus on 4 drug combinations: ceftriaxone (CRO) and ampicillin (AMP); ampicillin and streptomycin (STR); ceftriaxone and ciprofloxacin (CIP); and tigecycline (TGC) and ciprofloxacin. We describe these combinations in more detail below.

### Laboratory evolution across iso-inhibitory dosage combinations

Our goal was to compare evolutionary adaptation for different dosage combinations of each drug pair. The rate of adaptation is expected to depend heavily on the level of growth inhibition in the initial cultures, which sets the selection pressure favoring resistant mutants. To control for initial inhibition levels, we chose four dosage combinations for each drug pair—two corresponding to single drug treatments and two to drug combinations—that lie (approximately) along a contour of constant inhibition (i.e. constant per capita growth rate). We then evolved 24 replicate populations in each dosage combination for 3-4 days (20-30 generations) with daily dilutions into fresh media and drug. The evolution experiments were performed in microwell plates, allowing us to measure time series of cell density (OD) for each population over the course of the adaptation. In addition, we characterized the phenotypic resistance of 6 randomly chosen populations per condition by measuring standard dose-response curves and estimating the half-maximal inhibitory concentration (IC_50_) of each drug in the combination. Together, these measurements provide a detailed quantitative picture of both growth adaptation and changes in phenotypic resistance to each drug over the course of the evolution experiment.

### Dual β-lactam combination accelerates growth adaption but selects for similar resistance profiles as adaption to component drugs

Cell wall synthesis inhibitors, including β-lactams, are among the most frequently used antibiotics for *E. faecalis* infections (47). While *E. faecalis* often exhibit sensitivity to amino-pencillins, such as ampicillin (AMP), they are intrinsically resistant to cephalosporins (e.g. ceftriaxone (CRO)). Despite the limited utility of ceftriaxone alone, it combines with ampicillin to form a powerful synergistic pair, making it an attractive option for *E. faecalis* harboring high-level aminoglycoside resistance. Dual β-lactam combinations like CRO-AMP have been particularly effective in treatment of endocarditis infections in the clinical setting (44, 40).

As expected, we found that the CRO-AMP combination is strongly synergistic in the ancestral V583 *E. faecalis* strain (Figure 1a, left panel). We selected four dosage combinations (labeled A-D) along the concave contour of constant inhibition and evolved replicate populations to each condition. Growth curves on day 0, the first day of evolution, show similar levels of inhibition for each combination, but the day 2 curves vary substantially between replicates and between conditions (Figure 1a, right panels). To quantify growth, we estimated per capita growth rate during the early-exponential phase of growth for each day and each condition using nonlinear least squares fitting to an exponential function (Figure 1b, left panels; Figure S1). In all four conditions, growth increases significantly by day 2, though the rate of adaptation is considerably different across conditions. Notably, day 2 growth rate is higher when both drugs are present (conditions B and C) than it is for the single-drug conditions (A and B). To further quantify these trends, we estimated the average adaptation rate for each population using linear regression for each growth rate time series (Figure 1b, right panel). While the 24 replicate populations in each condition show considerable variability–as expected, perhaps, for a stochastic evolutionary process–growth adaptation is significantly faster for the two combinations (B and C) than for the single drug treatments (A and B).

**FIG 1.**
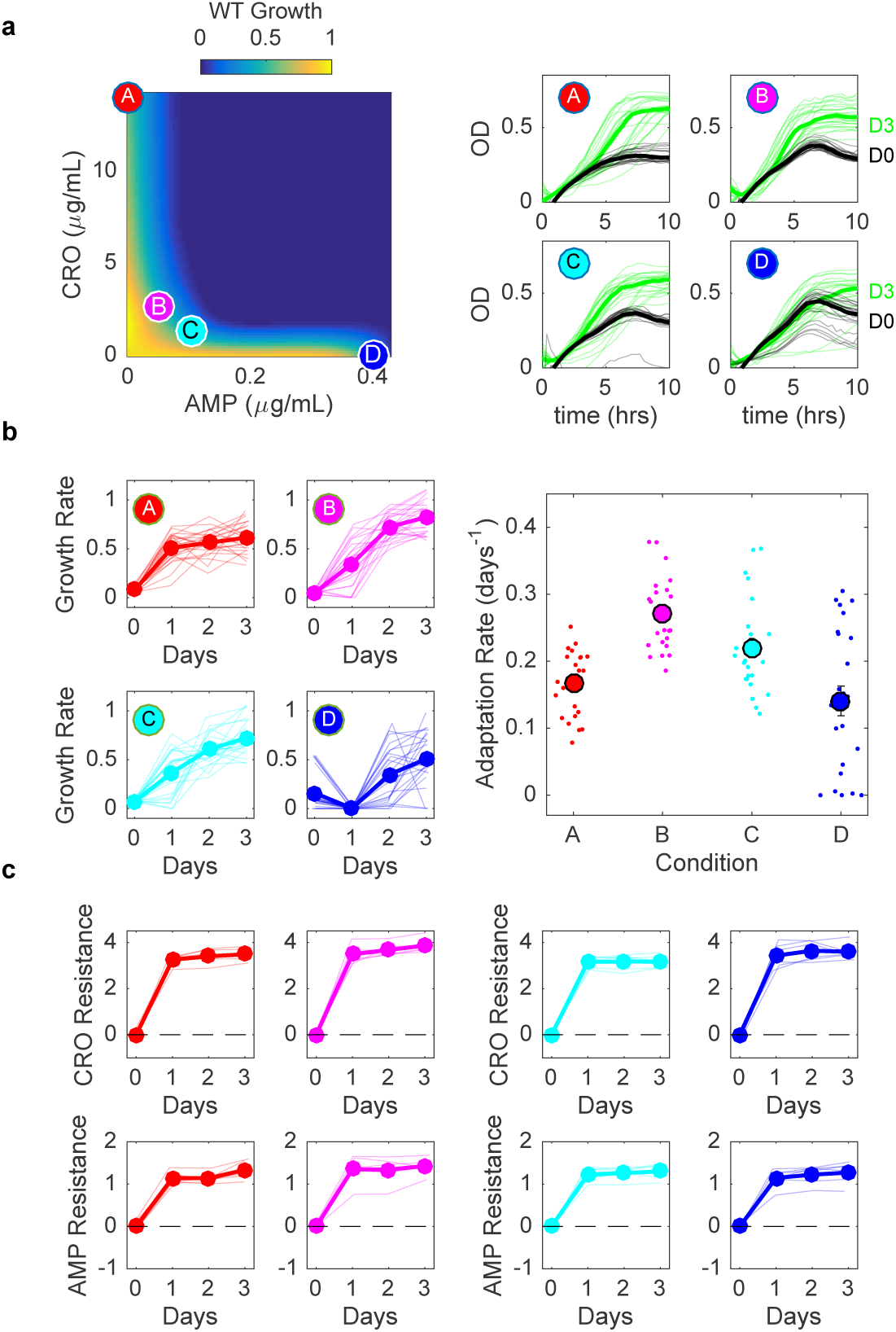
Dual β-lactam combination accelerates growth adaption but selects for similar resistance levels to component drugs. a) Left panel: per capita growth rate of ancestral populations as a function of ampicillin (AMP) and ceftriaxone (CRO) concentrations. Circles correspond to different selecting conditions along a contour of constant inhibition. Drug concentrations (AMP,CRO) are (0, 48) (red, A); (0.05, 2.6) (magenta, B); (0.10, 1.3) (cyan, C); and (0.43, 0) (blue, D). The latter point is shifted for visualization. Right panels: growth curves for the first (black) and last (green) days of evolution for each condition. b) Left panels: per capita growth rate over time for each condition. Right panel: average rate of growth adaptation over the course of the evolution. c. Resistance to CRO (top panels) and AMP (bottom panels) over time for isolates from different conditions. Resistance is defined as the log_2_-scaled fold change in IC_50_ of the resistant isolate relative to ancestral cells (positive is increased resistance, negative is increased sensitivity). In all plots, light transparent lines correspond to individual populations and darker lines to the mean across populations.

One might naively expect that increased growth adaption in the drug combination indicates that populations in these conditions evolve higher levels of resistance to one or both drugs. To test this hypothesis, we used replicate dose-response measurements to estimate the half-maximal inhibitory concentration (IC_50_) of both CRO and AMP in 6 randomly selected populations for each condition. We then quantified resistance as the (log_2_-scaled) fold change in IC_50_ between the evolved and ancestral populations; positive values indicate increased resistance and negative values indicate increased sensitivity relative to the ancestral strain (Figure 1c). Surprisingly, populations evolved under all four conditions exhibit strikingly similar patterns of phenotypic resistance, with IC_50_’s to each drug rising rapidly and plateauing after 1-2 days. In all conditions, populations tend to show higher increases in CRO resistance than AMP resistance. These results are initially surprising because they indicate that mutants with nearly identical phenotypic resistance profiles nevertheless exhibit markedly different patterns of growth adaptation that depend on the specific dosage combination.

### Aminoglycoside/β-lactam and β-lactam/fluoroquinolone combinations slow growth adaptation and select for resistant profiles distinct from those evolved to the component drugs

In addition to dual β-lactam therapies, combinations involving an aminoglycoside with a cell wall inhibiting antibiotic are commonly used for treating drug resistant *E. faecalis* (40). In particular, the ampicillin and streptomycin (STR) combination has been used as a first line of treatment for *E. faecalis* infective carditis (47, 40). Unfortunately, enterococci isolates are increasingly exhibiting high-level resistance to aminoglycosides, which has been shown to reduce the synergistic effect of the combination therapy (47). While aminoglycoside resistance is a growing problem, the adaptation of aminoglycoside-resistant *E. faecalis* to combination therapies remains poorly understood.

Consistent with previous findings, we did not observe synergy between AMP and STR in the ancestral V583 strain, which exhibits considerable aminoglycoside resistance. In fact, the drug pair exhibits marked antagonism, as evidenced by the convex growth contours on the response surface (Figure 2a, left panel). Growth curves from populations evolved to four different conditions along a growth contour show considerable differences at day 2 between the single-drug conditions (conditions A and D) and the two-drug conditions (B and C, Figure 2a, right panel; Figure S2), with single-drug populations reaching a higher growth rate (Figure 2b, left panels) and a dramatically increased rate of adaptation (Figure 2a, right panel). At the end of the experiment (day 3), populations grown in single drugs tend to show resistance to the selecting drug but little cross resistance to the other drug (Figure 2c, red and blue curves). Populations grown in a combination of both drugs show similar resistance profiles to those selected by the single drug that is dominant within the mixture. For example, the day 3 resistance profiles for condition B are similar to those from condition A (dominated by AMP), while those for condition C are similar to condition D (dominated by STR). It is interesting to note that the temporal dynamics (i.e. change in resistance over time for each drug) for all four conditions do show different qualitative trends, even when the final resistance profiles are similar (Figure 2c).

**FIG 2.**
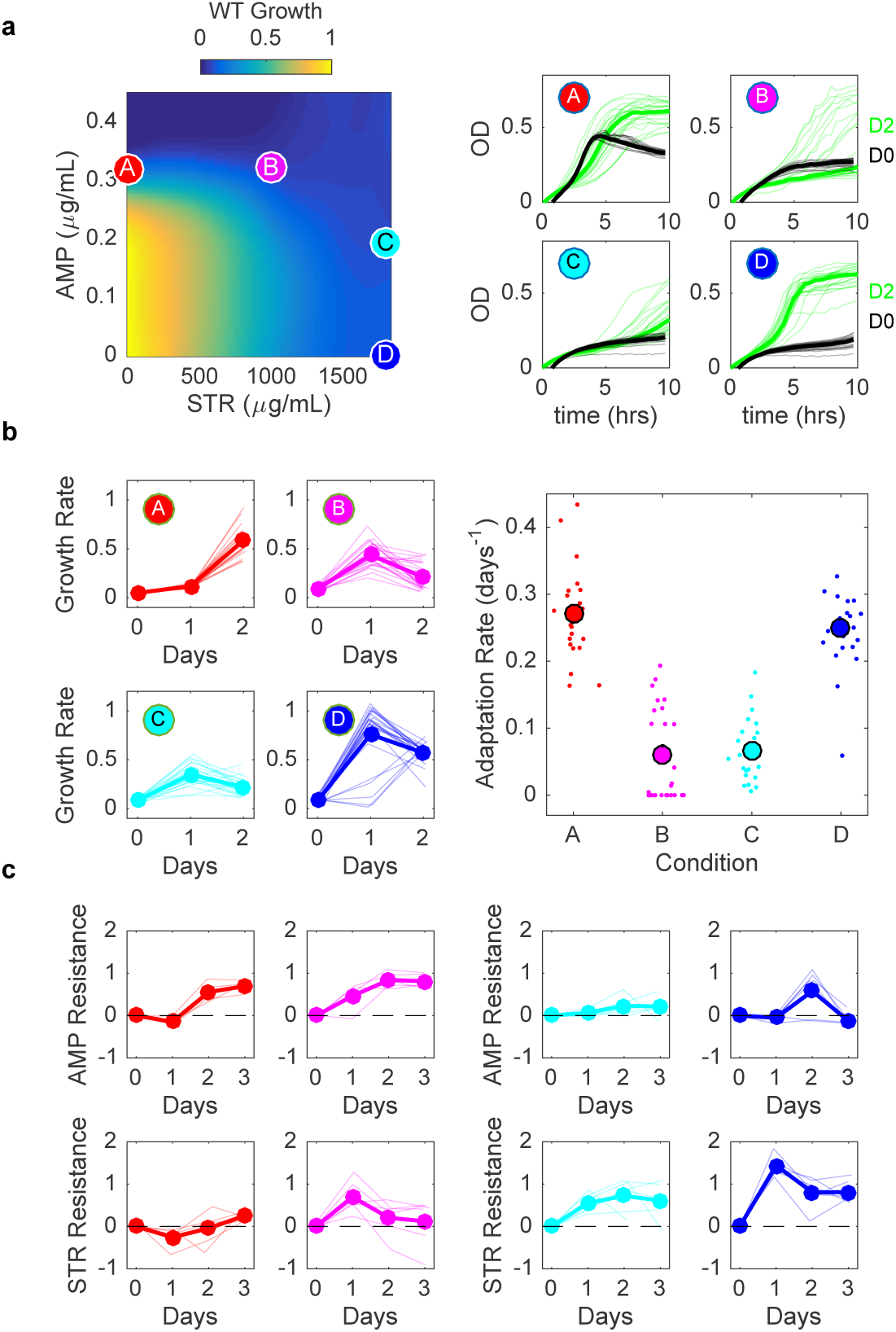
Antagonistic β-lactam and aminoglycoside combination slows adaptation with little cross-resistance. a) Left panel: per capita growth rate of ancestral populations as a function of ampicillin (AMP) and streptomycin (STR) concentrations. Circles correspond to different selecting conditions along a contour of constant inhibition. Drug concentrations (AMP,STR) are (0.3, 0) (red, A); (0.3, 1000) (magenta, B); (0.19, 1800) (cyan, C); and (0, 1800) (blue, D). Right panels: growth curves for the first (black) and last (green) days of evolution for each condition. b) Left panels: per capita growth rate over time for each condition. Right panel: average rate of growth adaptation over the course of the evolution. c. Resistance to AMP (top panels) and STR (bottom panels) over time for isolates from different conditions. Resistance is defined as the log_2_-scaled fold change in IC_50_ of the resistant isolate relative to ancestral cells (positive is increased resistance, negative is increased sensitivity). In all plots, light transparent lines correspond to individual populations and darker lines to the mean across populations

We observed qualitatively similar behavior in a combination of a β-lactam and fluoroquinolone (ciprofloxacin). Ciprofloxacin (CIP) is not typically used in the treatment of enterococci, though it has been used with β-lactams in the treatment of enterococcal endocarditis with high-level aminoglycoside resistance (48). In vitro studies also demonstrate efficacy of ciprofloxacin in multiple combinations against *E. faecalis* biofilms (49). We investigated resistance evolution in a combination of CIP with CRO, which is not a clinically used combination but exhibits less dramatic antagonism than the AMP-STR combination (Figure 3a, left panel), making it a potentially interesting proof-of-principle example of resistance involving fluoroquinolones.

**FIG 3.**
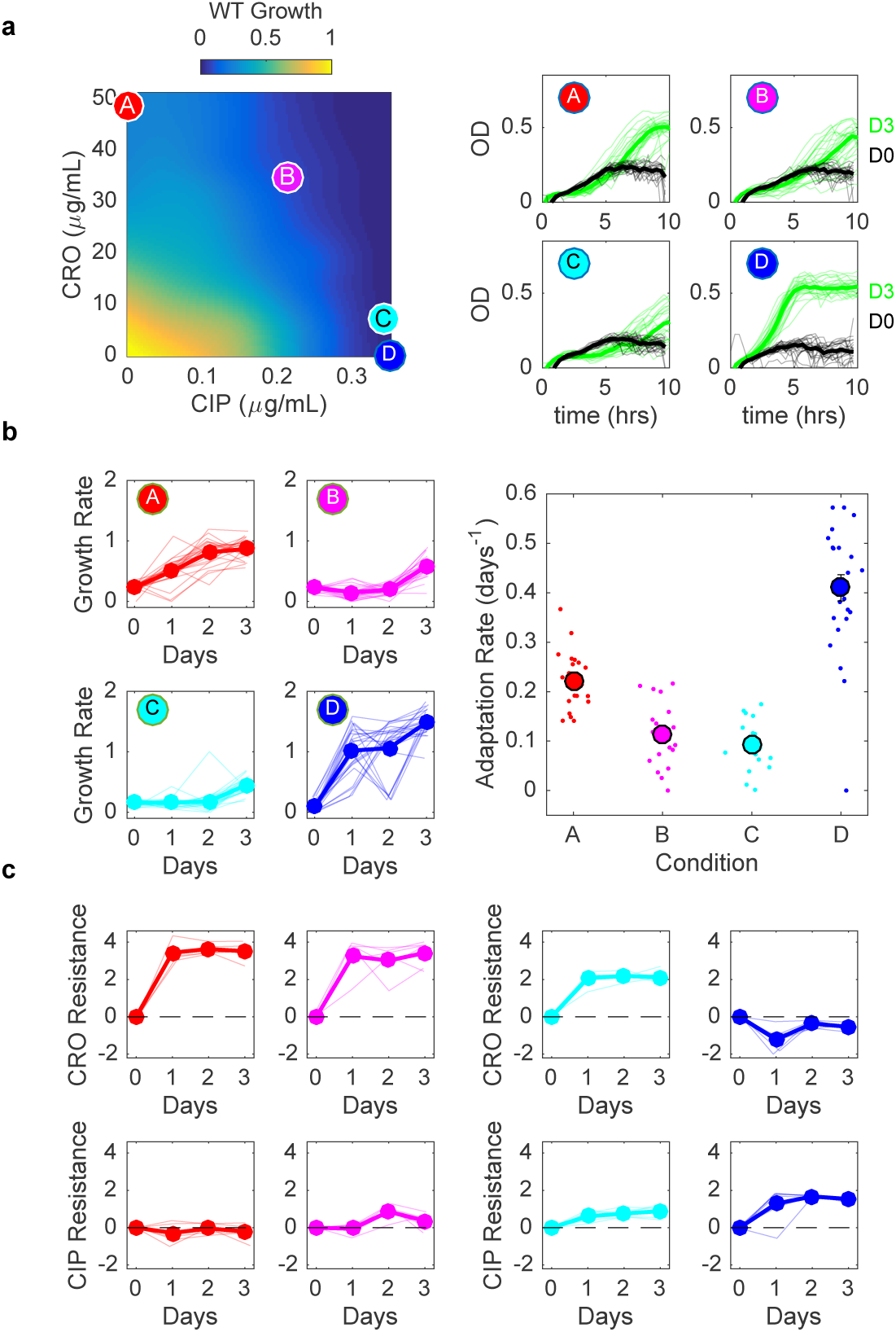
Antagonistic β-lactam and fluoroquinolone combination slows adaptation and selects for populations lacking observed collateral sensitivity of single-drug isolates. a) Left panel: per capita growth rate of ancestral populations as a function of ceftriaxone (CRO) and ciprofloxacin (CIP) concentrations. Circles correspond to different selecting conditions along a contour of constant inhibition. Drug concentrations (CRO, CIP) are (48.5, 0) (red, A); (34.7, 0.21) (magenta, B); (6.93, 0.34) (cyan, C); and (0, 0.46) (blue, D). The latter point is shifted for visualization. Right panels: growth curves for the first (black) and last (green) days of evolution for each condition. b) Left panels: per capita growth rate over time for each condition. Right panel: average rate of growth adaptation over the course of the evolution. c. Resistance to CRO (top panels) and CIP (bottom panels) over time for isolates from different conditions. Resistance is defined as the log_2_-scaled fold change in IC_50_ of the resistant isolate relative to ancestral cells (positive is increased resistance, negative is increased sensitivity). In all plots, light transparent lines correspond to individual populations and darker lines to the mean across populations.

As with the AMP-STR combination, we observe slowed growth adaptation in combinations of CRO-CIP relative to that in the component drugs alone (Figure 3b; Figure S3). Adaptation to CRO alone leads to strong CRO resistance and slight CIP sensitivity (Figure 3c, red). On the other hand, adaptation to CIP alone leads to CIP resistance along with significant increases in CRO sensitivity (Figure 3c, blue). Populations evolved to combinations show different resistance profiles at different concentrations, though for both mixtures the collateral sensitivities are eliminated (Figure 3c, magenta and cyan).

### Tigecycline suppresses growth adaptation and eliminates evolution of fluo-roquinolone resistance

The fourth combination we investigated was comprised of a protein synthesis inhibitor (tigecycline, TGC) and fluoroquinolone (CIP). TGC is a relatively new broad-spectrum antibiotic used for soft-tissue infections (50); it also shows in-vitro synergy in combination with multiple antibiotics (51, 52). We found that the TGC-CIP combination exhibits a particularly interesting type of interaction known as suppression (Figure 4a, left panel), where the combined effect of both drugs can be smaller than the effect of one drug (in this case, TGC) alone at the same concentration. Suppressive interactions between protein and DNA synthesis inhibitors in *E. coli* have been previously linked to sub-optimal regulation of ribosomal genes (53) as well as inverted selection for sensitive cells (37). Because the growth contours in this case show non-monotonic behavior, we performed evolution experiments in replicates of 8 for 11 different concentration combinations that fall along the isobole (Figure 4a; Figure S4). Strikingly, we find that growth adaptation decreases approximately monotonically as TGC concentration is increased, eventually approaching a minimum as TGC eclipses a critical concentration TGC*_crit_* ≈ 0.03 μg/mL (Figure 4c). Furthermore, while we observed TGC resistance only in rare cases, populations adapted to TGC below the critical concentration show approximately constant levels of CIP resistance, while those at higher concentrations show essentially no CIP resistance (Figure 4d). It is particularly striking that populations evolved in conditions A (red) and I (light blue) exhibit such different evolutionary behavior, as both are exposed to nearly identical CIP concentrations and, by design, start at similar levels of inhibition. Yet evolution in condition A leads to fast growth adaptation and strong CIP resistance, while evolution in condition I (light blue) leads to little adaptation and no CIP resistance. In effect, the addition of TGC eliminates CIP resistance without modulating the overall efficacy of the (initial) combination.

**FIG 4.**
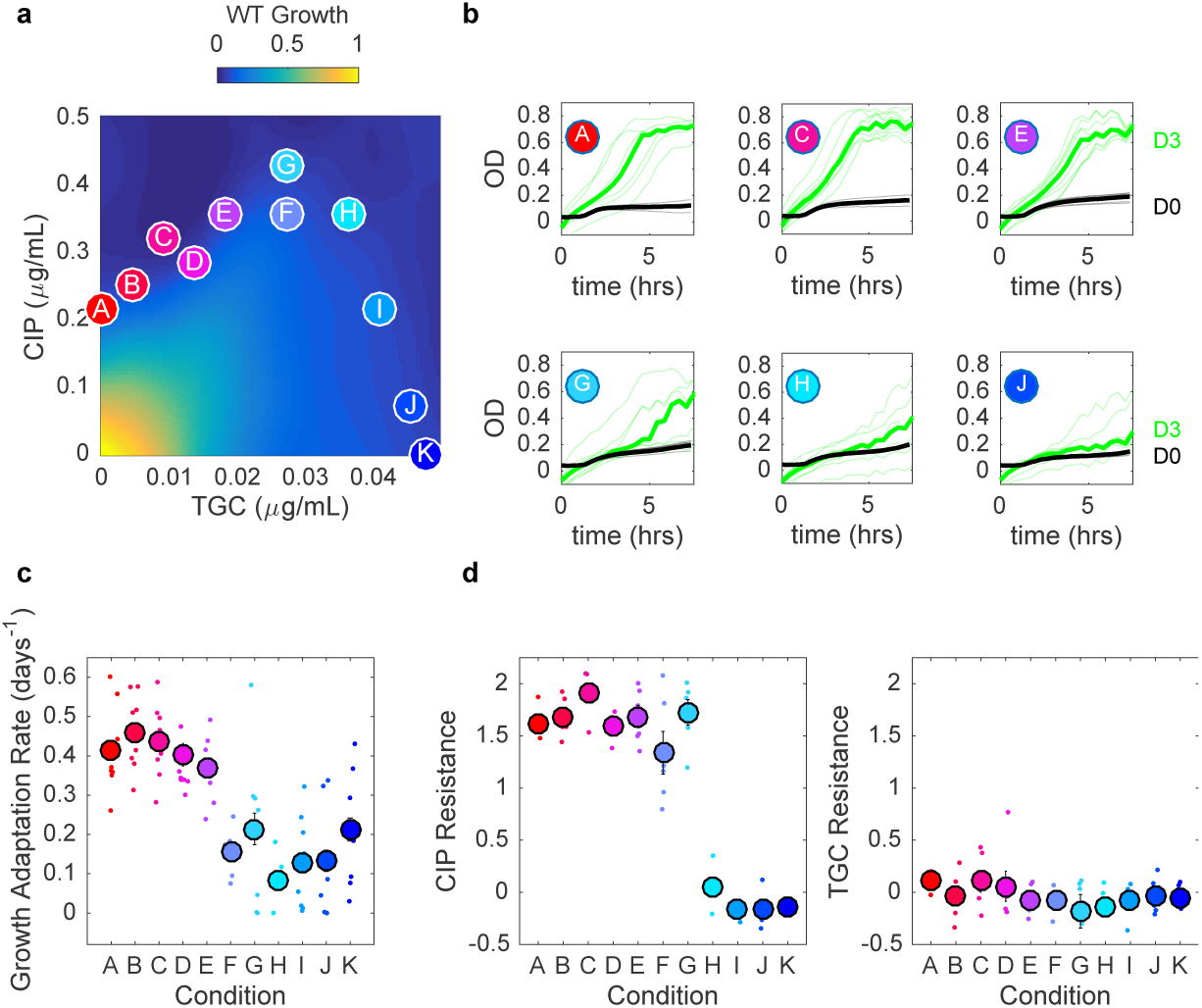
Tigecycline eliminates fluoroquinolone resistance above a critical concentration. a) Left panel: per capita growth rate of ancestral populations as a function of tigecycline (TGC) and ciprofloxacin (CIP) concentrations. Circles correspond to different selecting conditions along a contour of constant inhibition. b) growth curves for the first (black) and last (green) days of evolution for six of the 11 selecting conditions. c) Average rate of growth adaptation over the course of the evolution for each condition. d. Resistance to CIP (left) and TGC (right) for isolates on the final day of evolution. Resistance is defined as the log_2_-scaled fold change in IC_50_ of the resistant isolate relative to ancestral cells (positive is increased resistance, negative is increased sensitivity). In all plots, light transparent lines or small points correspond to individual populations. Darker lines and larger circles represent means taken across populations.

### Geometric rescaling of ancestral growth surface explains condition-dependent growth adaptation when resistant profiles are unchanged

To interpret the observed evolutionary dynamics, we hypothesized that mutations conferring resistance modulate the effective drug concentration experienced by the population. In some cases–for example, resistance due to efflux pumps–this effective concentration change corresponds to a genuine physical change in intracellular drug concentration. More generally, though, this hypothesis assumes that resistant cells growing in external drug concentration C behave similarly to wild type (drug-sensitive) cells experiencing a reduced effective concentration C’ < C. Similar rescaling arguments were pioneered in (35, 37), where they were used to predict correlations between the rate of resistance evolution and the type of drug interaction.

Our results indicate that adaptation rates in CRO-AMP (Figure 1) and TGC-CIP (Figure 4) combinations can vary significantly across dosage combinations, even when resistance profiles are essentially unchanged. For example, populations adapted to CRO-AMP combinations show an average increase in IC_50_ for CRO and AMP of about 2^3.5^ ≈ 11.6 fold and 2^1.3^ ≈ 2.5 fold, respectively. To understand how this level of resistance might be expected to impact growth, we rescaled the concentrations of CRO and AMP that lie along the contour of constant growth that passes through the four experimental dosage combinations (A-D). For each point on the contour, the concentrations of CRO and AMP are reduced by factors of 11.6 and 2.5, respectively. For example, the points corresponding to conditions A-D are mapped to the points shown in Figure 5a (squares). The new rescaled contour, which includes the rescaled locations of the original points A-D, does not in general correspond to a contour of constant growth on the original growth surface; therefore, growth of the adapted cells is expected to differ along the contour. More specifically, to predict growth of mutants selected along the original contour of constant growth, one simply needs to find the rescaled contour, plot it atop the original (ancestral) growth surface, and read off the values of growth along that rescaled contour. In the CRO-AMP combination, this rescaling approach predicts that growth is generally higher when the drugs are combined (e.g. conditions B and C) than when they are used individually (Figure 5a, top panel), in qualitative agreement with our experiments (Figure 1).

**FIG 5.**
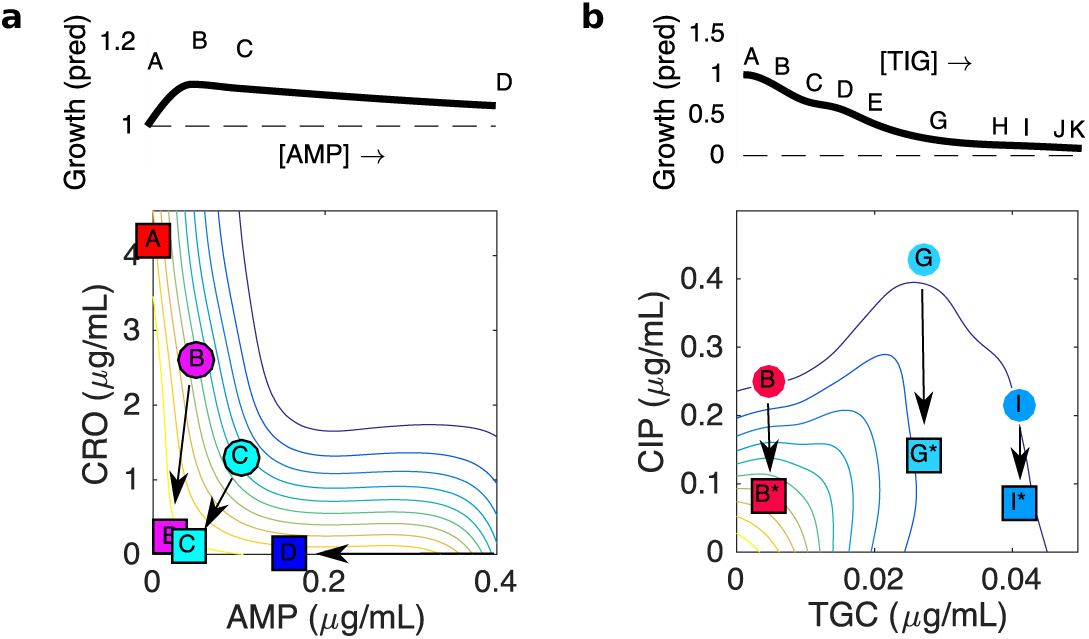
Geometric rescaling of growth surface in ancestral strain explains growth differences between different conditions even when resistance profiles are identical. Bottom panels, contour plots show growth in ancestral (WT) strain as function of drug concentrations for AMP-CRO (a, left) and TGC-CIP (b, right). Circles indicate selecting conditions (note that for visualization purposes, all four selecting conditions are not shown). Squares indicate effective concentrations achieved by rescaling true concentrations by the observed (mean) fold change in IC_50_ for each drug. All arrows in a single panel correspond to the same rescaling: AMP→ AMP/2.5, CRO→ CRO/11.6 (panel a) and CIP→ CIP/3.0, TGC → TCG (panel b). The rescaling factors correspond to cross resistance to AMP and CRO, with CRO resistance larger than AMP resistance (panel a) and to CIP resistance with no change in TGC resistance (panel b), which correspond to the mean values observed experimentally over all populations in AMP-CRO (see Figure 1) and the CIP-resistant populations in TGC-CIP (see Figure 4). Upper panels: predicted (relative) change in growth as one moves along the original contour, starting at condition A (one drug only). Predicted growth rate is calculated using 2d interpolation to estimate the growth along the rescaled contour on the ancestral growth surface.

Similarly, populations adapted in TGC-CIP show an increase in CIP IC_50_ of approximately 2^1.6^ ≈ 3 fold, but only when adaptation occurs below a critical TGC concentration. If we apply the same rescaling approach–that is, we reduce the concentration of CIP by 3-fold for all points along the contour–we again get a series of new points that no longer fall on a single growth contour (Figure 5b, squares). Furthermore, the predicted growth for points on the new contour decreases monotonically with TGC concentration before plateauing near point G, near the critical concentration where experimental growth adaptation approaches its minimum value (Figure 5b). Intuitively, then, it becomes clear why selection for ciprofloxacin resistance is only favored below this critical concentration: for higher concentrations of TGC, the rescaled points fall very nearly on the same contour as the original point. That is, when the original contour becomes approximately vertical, rescaling the CIP concentration is no longer expected to increase growth (see, for example, point I).

### Resistance profiles selected in different AMP-STR and CRO-CIP combinations are (nearly) growth-optimized linear combinations of the profiles selected by component drugs

Rescaling arguments may also help us to understand why particular resistance profiles appear to be preferentially selected under different initial conditions, as we observed with AMP-STR and CRO-CIP. In both cases, the resistance profiles on the final day of the experiment fall approximately on a line segment in the two dimensional space describing resistance to each drug (Figure 6a and b, left panels; line segments are labeled with endpoints X and Y). Because these line segments (approximately) connect points corresponding to profiles from conditions A and D (the single-drug conditions), resistance profiles along this line are linear combinations of the resistance profiles for singe-drug conditions.

**FIG 6.**
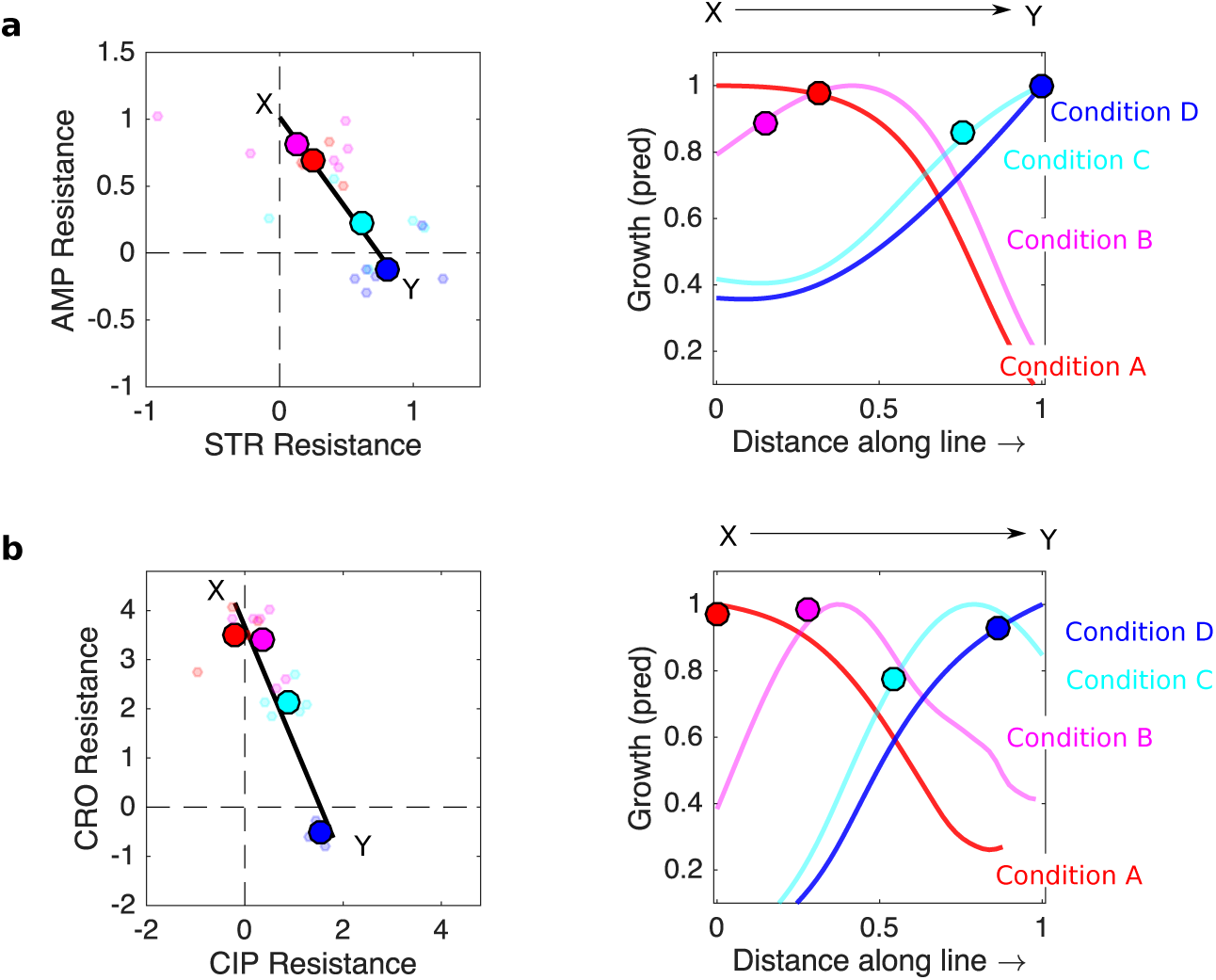
Resistance profiles observed in different AMP-STR and CRO-CIP dosage combinations are (nearly) growth-optimized linear combinations of profiles selected by component drugs. a: AMP-STR, b. CRO-CIP. Left panels: resistance profiles for individual populations (small circles) and the mean across populations (large filled circles) for each selecting condition. Color scheme is the same as in previous figures (i.e. red is condition A, magenta condition B, cyan condition C, and blue condition D). Solid black lines are linear fits to the averaged resistance profiles, which correspond (approximately) to the collection of profiles representing linear combinations (*r*_*lin*_ = *c*_1_*r*_1_ + (1 − *c*_1_)*r*_2_) of the profiles *r*_1_ and *r*_2_ selected by the component drugs. Right panels: each curve shows the predicted growth for populations with a range of resistance profiles (falling on the line XY) grown at one specific selecting condition (A, red; B, magenta; C, cyan; D, blue). The horizontal axis corresponds to position along the line segment XY (left panels). Filled circles correspond to the locations of experimentally observed (mean) resistance profiles.

To investigate how these linear combinations of single-drug profiles might be expected to affect growth in each selecting condition, we rescaled the drug concentrations corresponding to each selecting condition by a range of rescaling factors that lie along the line segment XY. As a result, each of the original selecting conditions (more specifically the points defined by the drug concentrations describing each condition) is mapped to a smooth curve in the two-drug concentration space. Points along that curve indicate how the original selecting concentrations are mapped to rescaled drug concentrations by the resistance profiles lying along segment XY. From that new curve, then, one can read off the corresponding growth rates on the ancestral growth surface, leading to predicted growth rates for mutants with any particular linear combination of the single drug profiles (i.e. any profile on line segment XY, Figure 6 a and b, right panels).

For each selecting condition, there is an optimal resistance profile (on the line segment XY) that leads to the maximum possible expected growth. Not surprisingly, in the case of AMP-STR, maximum growth in condition A (AMP only) occurs at point X, where AMP resistance is highest. Similarly, maximum growth in conditions D (STR only) and C occur at point Y, which has the largest STR resistance. On the other hand, the optimal resistance profile for condition B lies just short of the midpoint on the line segment XY (Figure 6a, right panel). Remarkably, the (mean) resistance profiles observed experimentally (circles) are predicted to give growth rates within approximately 15% of the optimal value.

In the case of the CRO-CIP combination, the optimal resistance profiles (of those that lie along line segment XY) for conditions A (CRO only) and D (CIP only) lie at the endpoints X and Y, respectively (Figure 6b, right panel), which have the highest resistance levels to the component drugs. By contrast, the optimal profiles for conditions B (magenta) and C (cyan) fall at different points along the XY segment, reflecting trade-offs between resistance levels and collateral sensitivities to the component drugs. Once again, the observed (mean) resistance profiles are predicted to give growth rates very near the optimal values for each condition (particularly for conditions A, B, and D). Given that there is a finite number of genetic mutations possible under these short-term conditions, one would not expect that phenotypic profiles exist for all points on the line segments XY. Nevertheless, these results suggest that for these two drug pairs, the resistant profiles selected in the combinations are linear combinations of the profiles selected by the component drugs alone. Even more surprisingly, the observed profiles are expected to give growth benefits that are nearly optimal among all the possible linear combinations.

## DISCUSSION

Using laboratory evolution, we have shown that adaptation of *E. faecalis* populations to drug combinations can differ substantially from adaptation to the component drugs. While the evolutionary trajectory of any particular population is difficult to predict, the results as a whole point to simple trends that can be explained with rescaling arguments linking growth of adapted populations to growth of the ancestral population at properly rescaled drug dosages. These arguments show, for example, how identical resistance profiles yield different growth rates for different selecting conditions. The analysis also suggests that, in multiple cases, the profiles selected by different dosage combinations are very nearly growth-optimized linear combinations of the profiles selected by the individual component drugs.

It is important to point out several limitations to our study. First, our goal was not to investigate the specific molecular mechanisms involved in drug adaption, but instead to provide a quantitative picture of resistance evolution that does not require extensive molecular-level knowledge, which many times is not available. However, the richness of the observed phenotypes points to complex and potentially interesting genetic changes that can be partially resolved with modern sequencing technologies. For example, cross-resistance observed between ceftriaxone and ampicillin may be due to mutations in penicillin-binding proteins, which are common resistance determinants for both drugs (47). We also note that our experiments were performed just below the minimum inhibitory drug concentrations, allowing for slowed but nonzero proliferation. Previous work indicates that drug interactions may modulate evolution in different ways at higher drug concentrations (27). In addition, our experiments were performed in planktonic populations, while many of the high-inoculum infections requiring combination treatment are likely to involve surface-associated biofilms, where spatial heterogeneity and complex community dynamics can dramatically alter the response to antibiotics. In *E. faecalis*, for example, population density can significantly modulate growth dynamics (8), while sub-inhibitory doses of cell wall inhibitors may actually promote biofilm growth (54). Recent work in other bacterial species also shows that evolutionary adaptation may differ between biofilm and planktonic communities (55, 56). Investigating adaptation to drug combinations in these different regimes, both at clinically-relevant concentrations and in biofilms, remains an interesting avenue for future work.

There are also several notable technical limitations. It is clear that all growth curves are not purely exponential, and in fact the per capita growth rate can change with time. Our growth rate estimates should therefore be thought of as an effective growth rate that reduces the population dynamics each day to a single number. Similarly, the rescaling analysis assumes that the drug resistance in the population can be captured by a single IC_50_ (for each drug), essentially neglecting clonal interference in favor of a single dominant resistant phenotype. In addition, the rescaling analysis does not incorporate fitness cost (which we did not measure). Each of these limitations could potentially be overcome with significantly more experimental data–for example, OD measurements at shorter intervals would allow for time-dependent growth rate estimates within each daily period, while resistance phenotyping of individual isolates from each population could be used to evaluate population heterogeneity. Given the potential complexity of evolutionary trajectories, even in these simplified laboratory scenarios, it is remarkable that simple rescaling arguments can qualitatively capture the coarse-grained features we measured. Future studies that aim to overcome the technical limitations of this work may be able to further evaluate quantitative agreement between specific evolutionary trajectories and the predictions of rescaling.

Most importantly, we stress that our results are based on in vitro laboratory experiments, which provide a well-controlled but potentially artificial–and certainly simplified–environment for evolutionary selection. While in vitro studies form the basis for many pharmacological regimens, the ultimate success or failure of new therapies must be evaluated using in vivo model systems and, ultimately, controlled clinical trials. We hope the results presented here offer a provocative look at evolution of *E. faecalis* in multidrug environments, but it is clear that these findings are not directly transferable to the clinic. Our results do include some clinically relevant antibiotic combinations, though we also sought a wide range of drug interaction types, including multiple antagonistic combinations that are unlikely a priori choices for clinical use. It is notable, however, that these non-synergistic combinations often produced considerably slower growth adaptation, consistent with previous results that highlight potential benefits for non-standard combinations (34, 35, 37).

Finally, our results reveal that simple rescaling arguments—similar to those originally introduced in (35, 37)—can be used to understand many features of evolution in two-drug environments. Extending and formalizing these qualitative findings using stochastic models where fast evolutionary dynamics are coupled to geometric rescaling on adiabatically changing interaction landscapes is an exciting avenue for theoretical work that may provide both general insight as well as specific, experimentally testable predictions for how resistance evolves in multi-drug environments.

**TABLE 1.** Antibiotics Used in this study

| Antibiotic (abbrev.) | Class | Mechanism of Action |
| --- | --- | --- |
| Ampicillin (AMP) | β-lactam (penicillins) | cell wall synthesis inhibitor |
| Ceftriaxone (CRO) | β-lactam (cephalosporin) | cell wall synthesis inhibitor |
| Ciprofloxacin (CIP) | fluoroquinolone | DNA synthesis inhibitor |
| Streptomycin (STR) | aminoglycoside | protein synthesis inhibitor |
| Tigecycline (TGC) | glycylcycline | protein synthesis inhibitor |

## MATERIALS AND METHODS

### Strains, media, and growth conditions

All experiments were performed on the V583 strain of *E. faecalis*, a fully sequenced clinical isolate. Overnight seed cultures were inoculated from a single colony and grown in sterilized brain heart infusion (BHI) medium at 37C with no shaking. Antibiotics stock solutions were prepared using sterilized Millipore water, diluted and aliquoted into single use micro-centrifuge tubes and stored at −20C or −80C. All drugs and media were purchased from Dot Scientific, Sigma-Aldrich or Fisher Scientific.

### Laboratory evolution and growth measurement

Evolution experiments were seeded by diluting overnight cultures of ancestral V583 cells 400x into individual wells of a 96-well microplate containing appropriate drug concentrations. All plates for a multi-day evolution experiment were prepared in advance by adding appropriate drug concentrations to 200 *μ*L BHI and storing at −20C for not more than 4 days. Each day, a new plate was thawed and inoculated with 2 *μ*L (100X dilution) from the previous day’s culture. Plates were then sealed with BIO-RAD Microseal film to minimize evaporation and prevent cross contamination. Optical density at 600 nm (OD) was measured for each population every 20-25 minutes using an EnSpire Multimode Plate Reader with multi-plate stacker attachment located in a temperature-controlled (30C) warm room. Control wells containing ancestral cells and BHI medium were included on each plate as a growth control and for background subtraction, respectively. All dilutions and daily transfers were performed inside a ThermoFisher 1300 Series A2 safety cabinet to minimize contamination. Samples from each population were stocked in 15 percent glycerol and stored at −80C.

### Estimating per capita growth rate and drug response surfaces

We estimated per capita growth rate from OD time series by fitting the early exponential phase portion of the background subtracted curves (typically OD<0.4) to an exponential function using nonlinear least squares (MATLAB 7.6.0 curve fitting toolbox, Mathworks). We normalized all growth rates by the growth rate of ancestral cells in the absence of drugs. We visualized two-drug growth response surfaces by smoothing (2-d cubic spline interpolation) to reduce experimental noise and displaying smoothed surfaces as two-dimensional heat maps. When relevant, growth at unsampled regions of the growth surface was estimated with 2d interpolation.

### Phenotypic resistance profiling

Experiments to estimate the half-maximal inhibitory concentration (IC_50_) for each population were performed in replicates of 4-8 in 96-well plates. Prior to IC_50_ testing, frozen stocks for each population were swabbed and grown overnight in drug-free medium. These overnight cultures were then diluted 100X into new plates containing fresh media and a gradient of 6-14 drug concentrations. After 20 hours of growth the optical density at 600 nm (OD600) was again measured and used to create a dose response curve. To quantify drug resistance, the resulting dose response curve was fit to a Hill-like function *f* (*x*) = (1+ (*x* /*K*) *^h^*)^-1^ using nonlinear least squares fitting, where *K* is the half-maximal inhibitory concentration (*IC*_50_) and *h* is a Hill coefficient describing the steepness of the dose-response relationship.

## ACKNOWLEDGMENTS

This work is supported by the National Science Foundation (NSF No. 1553028 to KW) and the National Institutes of Health (NIH No. 1R35GM124875-01 to KW). The format for this preprint is adapted from the American Society for Microbiology (ASM) template available on Overleaf.com.

## SUPPLEMENTAL MATERIAL

The Supplemental Material contains 4 supplemental figures (S1-S4).

---

**FIG S1.**
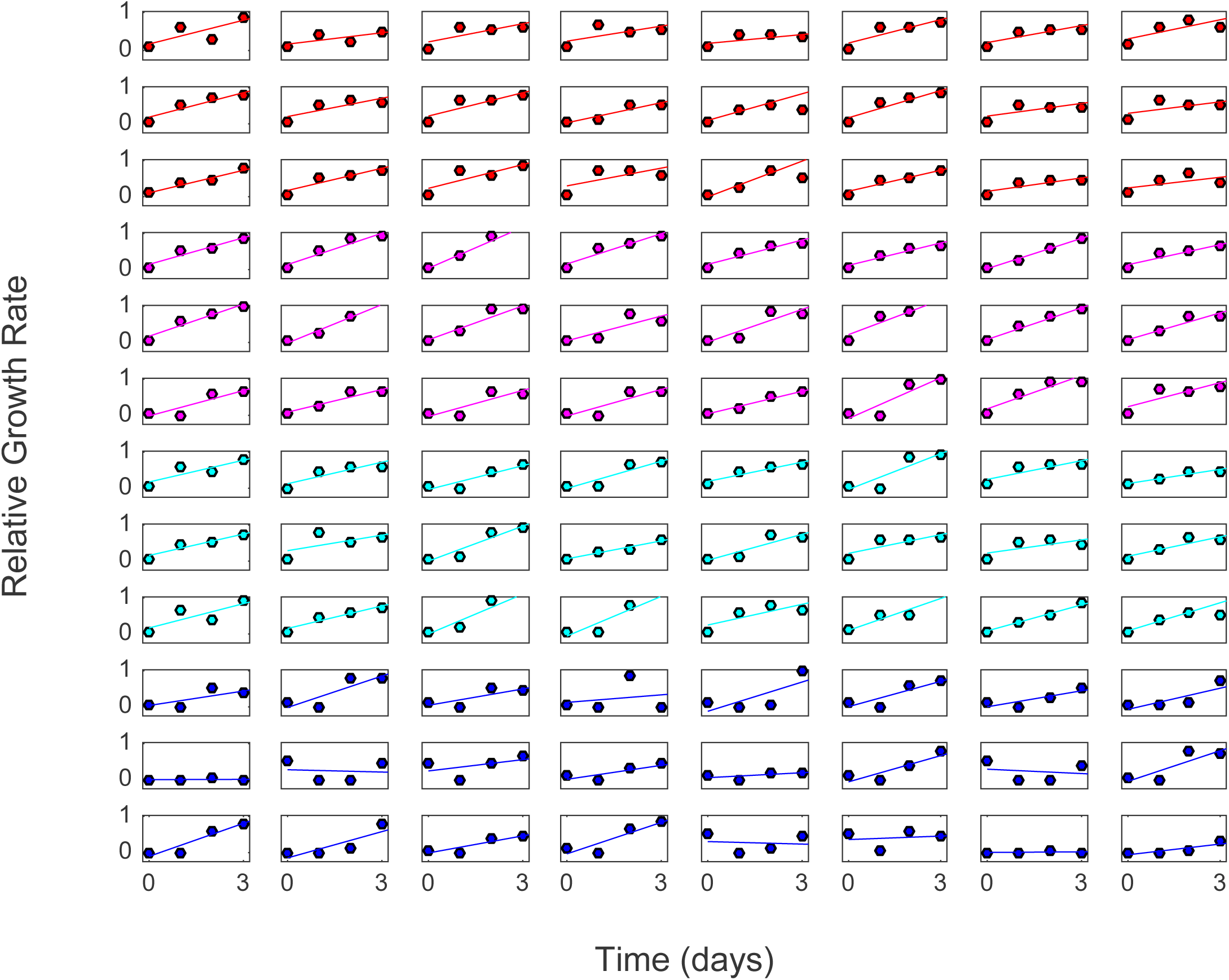
Growth rate time series (circles) and linear fits to determine mean adaptation rate (lines) for populations grown in conditions A (top 3 rows, red), B (magenta), C (cyan), and D (last 3 rows, blue) for combinations of ceftriaxone (CRO) and ampicillin (AMP).

**FIG S2.**
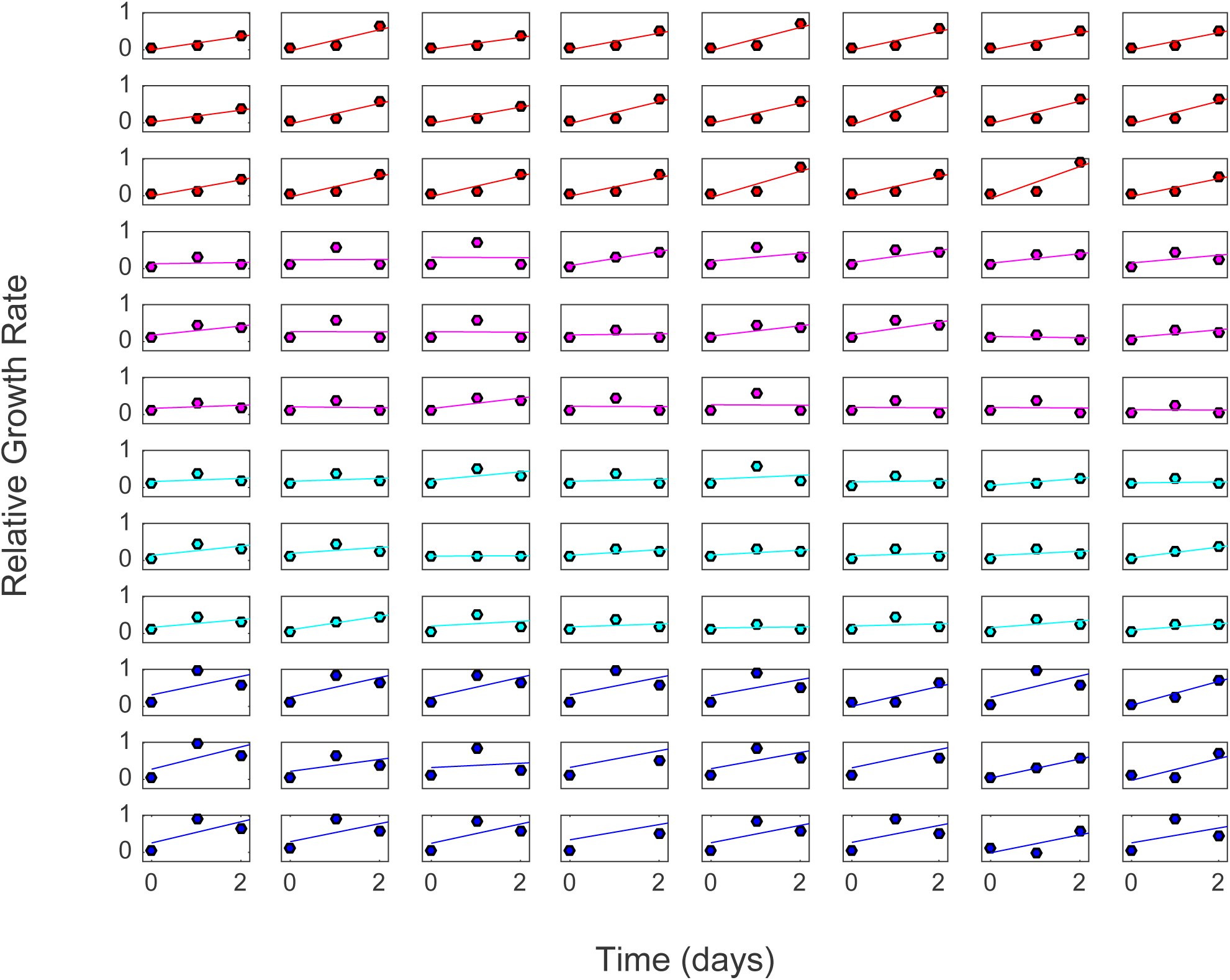
Growth rate time series (circles) and linear fits to determine mean adaptation rate (lines) for populations grown in conditions A (top 3 rows, red), B (magenta), C (cyan), and D (last 3 rows, blue) for combinations of streptomycin (STR) and ampicillin (AMP).

**FIG S3.**
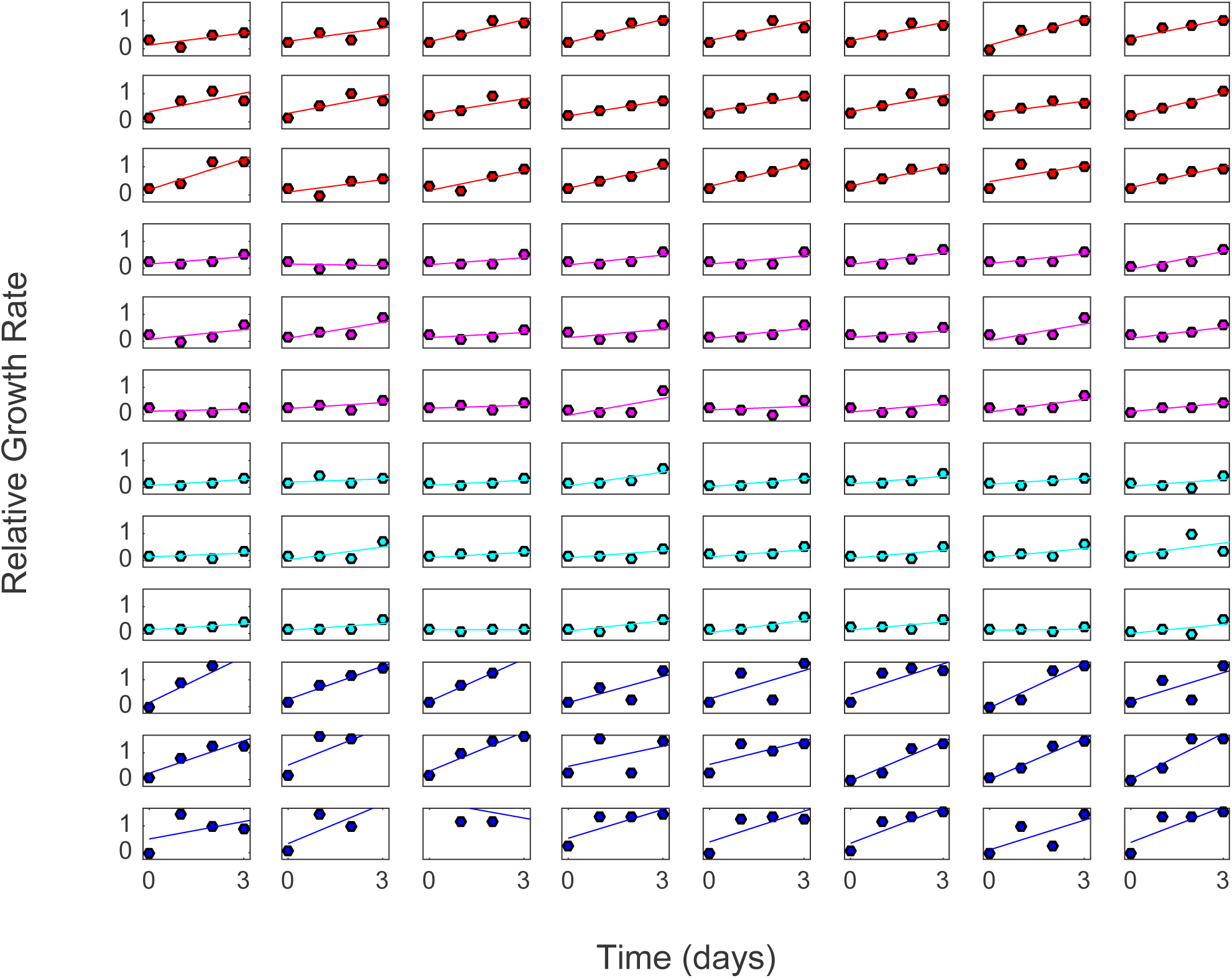
Growth rate time series (circles) and linear fits to determine mean adaptation rate (lines) for populations grown in conditions A (top 3 rows, red), B (magenta), C (cyan), and D (last 3 rows, blue) for combinations of ceftriaxone (CRO) and ciprofloxacin (CIP).

**FIG S4.**
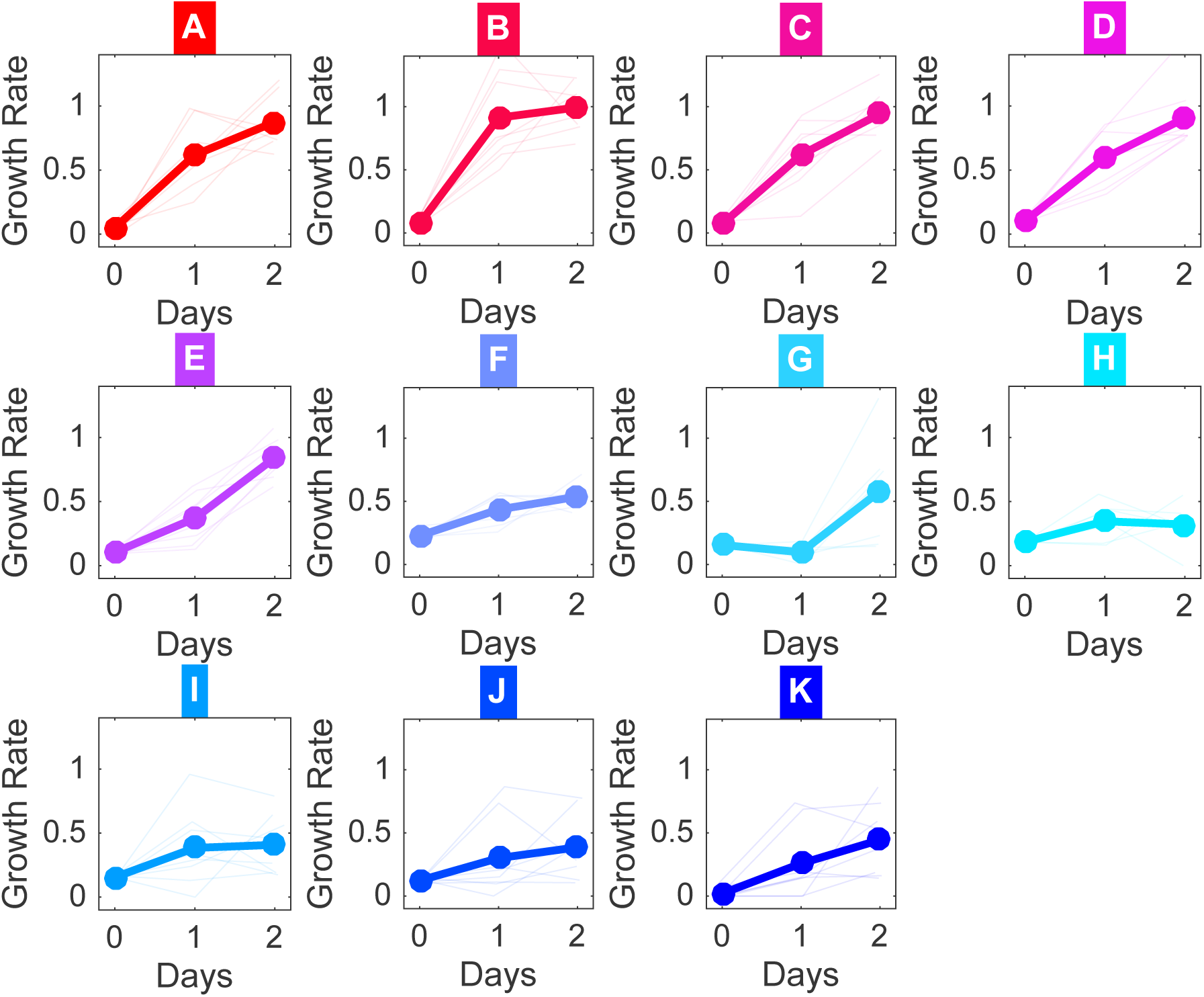
Growth rate time series for each population (light lines) and the mean across populations for a given condition for combinations of tigecycline (TGC) and ciprofloxacin (CIP).

## References

1. Davies J, Davies D. Origins and evolution of antibiotic resistance. Microbiol. Mol. Biol. Rev. 2010; 74(3):417–433.

2. Levy SB, Marshall B. Antibacterial resistance worldwide: causes, challenges and responses. Nat. medicine 2004; 10(12s):S122.

3. Read AF, Day T, Huijben S. The evolution of drug resistance and the curious orthodoxy of aggressive chemotherapy. Proc. Natl. Acad. Sci. 2011; 108(Supplement 2):10871–10877.

4. Hansen E, Woods RJ, Read AF. How to use a chemotherapeutic agent when resistance to it threatens the patient. PLoS biology 2017; 15(2):e2001110.

5. Meredith HR, Srimani JK, Lee A J, Lopatkin A J, You L. Collective antibiotic tolerance: mechanisms, dynamics and intervention. Nat. chemical biology 2015; 11(3):182–188.

6. Sorg RA, Lin L, Van Doorn GS, Sorg M, Olson J, Nizet V, Veening JW. Collective resistance in microbial communities by intracellular antibiotic deactivation. PLoS biology 2016; 14(12):e2000631.

7. Tan C, Smith RP, Srimani JK, Riccione KA, Prasada S, Kuehn M, You L. The inoculum effect and band-pass bacterial response to periodic antibiotic treatment. Mol. systems biology 2012; 8(1):617.

8. Karslake J, Maltas J, Brumm P, Wood KB. Population density modulates drug inhibition and gives rise to potential bistability of treatment outcomes for bacterial infections. PLoS computational biology 2016; 12(10):e1005098.

9. Gjini E, Brito PH. Integrating antimicrobial therapy with host immunity to 1ght drug-resistant infections: classical vs. adaptive treatment. PLoS computational biology 2016; 12(4):e1004857.

10. Zhang Q, Lambert G, Liao D, Kim H, Robin K, Tung Ck, Pourmand N, Austin RH. Acceleration of emergence of bacterial antibiotic resistance in connected microenvironments. Science 2011; 333(6050):1764–1767.

11. Baym M, Lieberman TD, Kelsic ED, Chait R, Gross R, Yelin I, Kishony R. Spatiotemporal microbial evolution on antibiotic landscapes. Science 2016; 353(6304):1147–1151.

12. Greulich P, Waclaw B, Allen RJ. Mutational pathway determines whether drug gradients accelerate evolution of drug-resistant cells. Phys. Rev. Lett. 2012; 109(8):088101.

13. Hermsen R, Deris JB, Hwa T. On the rapidity of antibiotic resistance evolution facilitated by a concentration gradient. Proc. Natl. Acad. Sci. 2012; 109(27):10775–10780.

14. Moreno-Gamez S, Hill AL, Rosenbloom DI, Petrov DA, Nowak MA, Pennings PS. Imperfect drug penetration leads to spatial monotherapy and rapid evolution of multidrug resistance. Proc. Natl. Acad. Sci. 2015; 112(22):E2874–E2883.

15. De Jong MG, Wood KB. Tuning spatial profiles of selection pressure to modulate the evolution of drug resistance. Phys. review letters 2018; 120(23):238102.

16. Trindade S, Sousa A, Xavier KB, Dionisio F, Ferreira MG, Gordo I. Positive epistasis drives the acquisition of multidrug resistance. PLoS genetics 2009; 5(7):e1000578.

17. Borrell S, Teo Y, Giardina F, Streicher EM, Klopper M, Feldmann J, Müller B, Victor TC, Gagneux S. Epistasis between antibiotic resistance mutations drives the evolution of extensively drug-resistant tuberculosis. Evol. medicine, public health 2013; 2013(1):65–74.

18. Yoshida M, Reyes SG, Tsudo S, Horinouchi T, Furusawa C, Cronin L. Time-programmable dosing allows the manipulation, suppression and reversal of antibiotic drug resistance *in vitro*. Nat. Commun. 2017; 8.

19. Meredith HR, Lopatkin A J, Anderson DJ, You L. Bacterial temporal dynamics enable optimal design of antibiotic treatment. PLoS computational biology 2015; 11(4):e1004201.

20. Nichol D, Jeavons P, Fletcher AG, Bonomo RA, Maini PK, Paul JL, Gatenby RA, Anderson AR, Scott JG. Steering evolution with sequential therapy to prevent the emergence of bacterial antibiotic resistance. PLoS computational biology 2015; 11(9):e1004493.

21. Fuentes-Hernandez A, Plucain J, Gori F, Pena-Miller R, Reding C, Jansen G, Schulenburg H, Gudelj I, Beardmore R. Using a sequential regimen to eliminate bacteria at sublethal antibiotic dosages. PLoS biology 2015; 13(4):e1002104.

22. Imamovic L, Sommer MOA. Use of collateral sensitivity networks to design drug cycling protocols that avoid resistance development. Sci. Transl. Med 2013; 5:204ra132.

23. Kim S, Lieberman TD, Kishony R. Alternating antibiotic treatments constrain evolutionary paths to multidrug resistance. Proc. Natl. Acad. Sci. USA 2014; 111:14494–14499.

24. Pál C, Papp B, Lázár V. Collateral sensitivity of antibiotic-resistant microbes. Trends microbiology 2015; 23(7):401–407.

25. Barbosa C, Trebosc V, Kemmer C, Rosenstiel P, Beardmore R, Schulenburg H, Jansen G. Alternative evolutionary paths to bacterial antibiotic resistance cause distinct collateral effects. Mol. biology evolution 2017; 34(9):2229–2244.

26. Barbosa C, Beardmore R, Schulenburg H, Jansen G. Antibiotic combination effcacy (ACE) networks for a Pseudomonas aeruginosa model. PLoS biology 2018; 16(4):e2004356.

27. Rodriguez de Evgrafov M, Gumpert H, Munck C, Thomsen TT, Sommer MO. Collateral resistance and sensitivity modulate evolution of high-level resistance to drug combination treatment in Staphylococcus aureus. Mol. biology evolution 2015; 32(5):1175–1185.

28. Nichol D, Rutter J, Bryant C, Hujer AM, Lek S, Adams MD, Jeavons P, Anderson AR, Bonomo RA, Scott JG. Antibiotic collateral sensitivity is contingent on the repeatability of evolution. Nat. communications 2019; 10(1):334.

29. Maltas J, Wood KB. Pervasive and diverse collateral sensitivity profiles inform optimal strategies to limit antibiotic resistance. bioRxiv 2019; https://www.biorxiv.org/content/early/2019/01/04/241075.

30. Podnecky NL, Fredheim EGA, Kloos J, Sorum V, Primicerio R, Roberts AP, Rozen DE, Samuelsen O, Johnsen PJ. Conserved collateral antibiotic susceptibility networks in diverse clinical strains of Escherichia coli. Nat. Commun. 2018; 9.

31. Imamovic L, Ellabaan MMH, Machado AMD, Citterio L, Wulff T, Molin S, Johansen HK, Sommer MOA. Drug-driven phenotypic convergence supports rational treatment strategies of chronic infections. Cell 2018; 172(1-2):121–134.

32. Baym M, Stone LK, Kishony R. Multidrug evolutionary strategies to reverse antibiotic resistance. Science 2016; 351(6268):aad3292.

33. Greco WR, Bravo G, Parsons JC. The search for synergy: a critical review from a response surface perspective. Pharmacol. reviews 1995; 47(2):331–385.

34. Michel JB, Yeh PJ, Chait R, Moellering RC, Kishony R. Drug interactions modulate the potential for evolution of resistance. Proc. Natl. Acad. Sci. 2008; 105(39):14918–14923.

35. Hegreness M, Shoresh N, Damian D, Hartl D, Kishony R. Accelerated evolution of resistance in multidrug environments. Proc. Natl. Acad. Sci. 2008; 105(37):13977–13981.

36. Pena-Miller R, Laehnemann D, Jansen G, Fuentes-Hernandez A, Rosenstiel P, Schulenburg H, Beardmore R. When the most potent combination of antibiotics selects for the greatest bacterial load: the smile-frown transition. PLoS biology 2013; 11(4):e1001540.

37. Chait R, Craney A, Kishony R. Antibiotic interactions that select against resistance. Nature 2007; 446(7136):668.

38. Torella JP, Chait R, Kishony R. Optimal drug synergy in antimicrobial treatments. PLoS computational biology 2010; 6(6):e1000796.

39. Munck C, Gumpert HK, Wallin AIN, Wang HH, Sommer MO. Prediction of resistance development against drug combinations by collateral responses to component drugs. Sci. translational medicine 2014; 6(262):262ra156–262ra156.

40. Beganovic M, Luther MK, Rice LB, Arias CA, Rybak MJ, LaPlante KL. A review of combination antimicrobial therapy for Enterococcus faecalis bloodstream infections and infective endocarditis. Clin. Infect. Dis. 2018; 67(2):303–309.

41. Clewell DB, Gilmore MS, Ike Y, Shankar N. Enterococci: from commensals to leading causes of drug resistant infection. Massachusetts Eye and Ear In1rmary; 2014.

42. Baddour LM, Wilson WR, Bayer AS, Fowler Jr VG, Tleyjeh IM, Rybak MJ, Barsic B, Lockhart PB, Gewitz MH, Levison ME, et al. Infective endocarditis in adults: diagnosis, antimicrobial therapy, and management of complications: a scienti1c statement for healthcare professionals from the American Heart Association. Circulation 2015; 132(15):1435–1486.

43. Chirouze C, Athan E, Alla F, Chu VH, Corey GR, Selton-Suty C, Erpelding ML, Miro JM, Olaison L, Hoen B, et al. Enterococcal endocarditis in the beginning of the 21st century: analysis from the International Collaboration on Endocarditis-Prospective Cohort Study. Clin. microbiology infection 2013; 19(12):1140–1147.

44. Fernández-Hidalgo N, Almirante B, Gavaldà J, Gurgui M, Peña C, De Alarcón A, Ruiz J, Vilacosta I, Montejo M, Vallejo N, et al. Ampicillin plus ceftriaxone is as effective as ampicillin plus gentamicin for treating Enterococcus faecalis infective endocarditis. Clin. infectious diseases 2013; 56(9):1261–1268.

45. Gavalda J, Len O, Miró JM, Munoz P, Montejo M, Alarcón A, De La Torre-Cisneros J, Pena C, Martínez-Lacasa X, Sarria C, et al. Brief communication: treatment of Enterococcus faecalis endocarditis with ampicillin plus ceftriaxone. Annals internal medicine 2007; 146(8):574–579.

46. Loewe S. The problem of synergism and antagonism of combined drugs. Arzneimittelforschung 1953; 3:285–290.

47. Kristich CJ, Rice LB, Arias CA. Enterococcal infection—treatment and antibiotic resistance. In: Enterococci: From commensals to leading causes of drug resistant infection [Internet] Massachusetts Eye and Ear In1rmary; 2014.

48. Tripodi M, Locatelli A, Adinolfi L, Andreana A, Utili R. Successful treatment with ampicillin and fluoroquinolones of human endocarditis due to high-level gentamicin-resistant enterococci. Eur. J. Clin. Microbiol. Infect. Dis. 1998; 17(10):734–736.

49. Holmberg A, Mörgelin M, Rasmussen M. Effectiveness of ciprofloxacin or linezolid in combination with rifampicin against Enterococcus faecalis in biofilms. J. antimicrobial chemotherapy 2011; 67(2):433–439.

50. Arias CA, Contreras GA, Murray BE. Management of multidrug-resistant enterococcal infections. Clin. microbiology infection 2010; 16(6):555–562.

51. Tang HJ, Chen CC, Zhang CC, Su BA, Li CM, Weng TC, Chiang SR, Ko WC, Chuang YC. In vitro effcacy of fosfomycin-based combinations against clinical vancomycin-resistant Enterococcus isolates. Diagn. microbiology infectious disease 2013; 77(3):254–257.

52. Silvestri C, Cirioni O, Arzeni D, Ghiselli R, Simonetti O, Orlando F, Ganzetti G, Staffolani S, Brescini L, Provinciali M, et al. In vitro activity and in vivo effcacy of tigecycline alone and in combination with daptomycin and rifampin against Gram-positive cocci isolated from surgical wound infection. Eur. journal clinical microbiology & infectious diseases 2012; 31(8):1759–1764.

53. Bollenbach T, Quan S, Chait R, Kishony R. Nonoptimal microbial response to antibiotics underlies suppressive drug interactions. Cell 2009; 139(4):707–718.

54. Yu W, Hallinen KM, Wood KB. Interplay between antibiotic effcacy and druginduced lysis underlies enhanced biofllm formation at subinhibitory drug concentrations. Antimicrob. agents chemotherapy 2018; 62(1):e01603–17.

55. Santos-Lopez A, Marshall CW, Scribner MR, Snyder D, Cooper VS. Biofilm-dependent evolutionary pathways to antibiotic resistance. bioRxiv 2019; p. 581611.

56. Martin M, Hölscher T, Dragoš A, Cooper VS, Kovács ÁT. Laboratory evolution of microbial interactions in bacterial biofilms. J. bacteriology 2016; 198(19):2564–2571.

