## Supplemental Text for "Antibiotic interactions shape short-term evolution of resistance in *E. faecalis*"

### SUPPLEMENTAL MATERIAL

The Supplemental Material contains 4 supplemental figures (S1-S4).

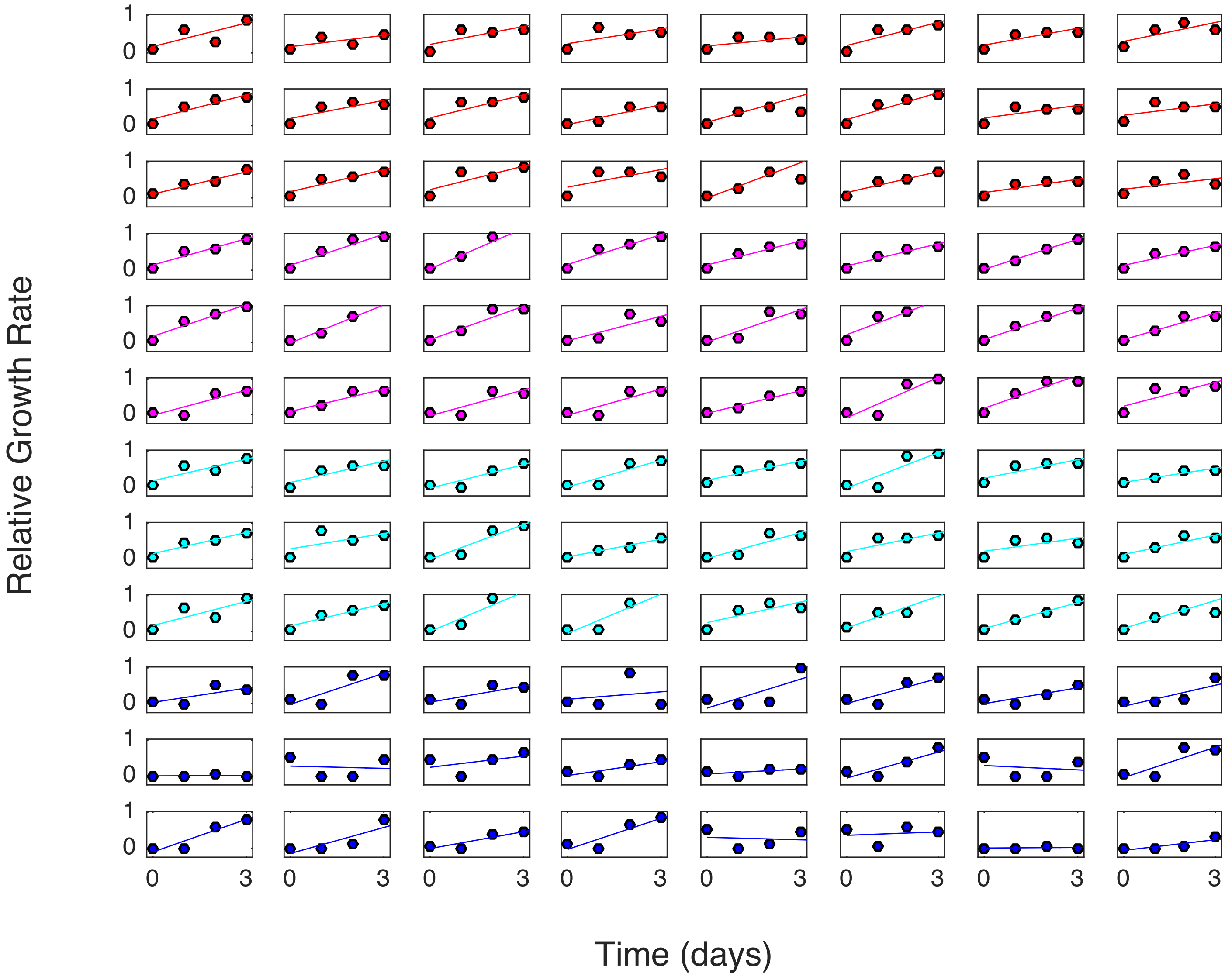


**FIG S1** Growth rate time series (circles) and linear fits to determine mean adaptation rate (lines) for populations grown in conditions A (top 3 rows, red), B (magenta), C (cyan), and D (last 3 rows, blue) for combinations of ceftriaxone (CRO) and ampicillin (AMP).

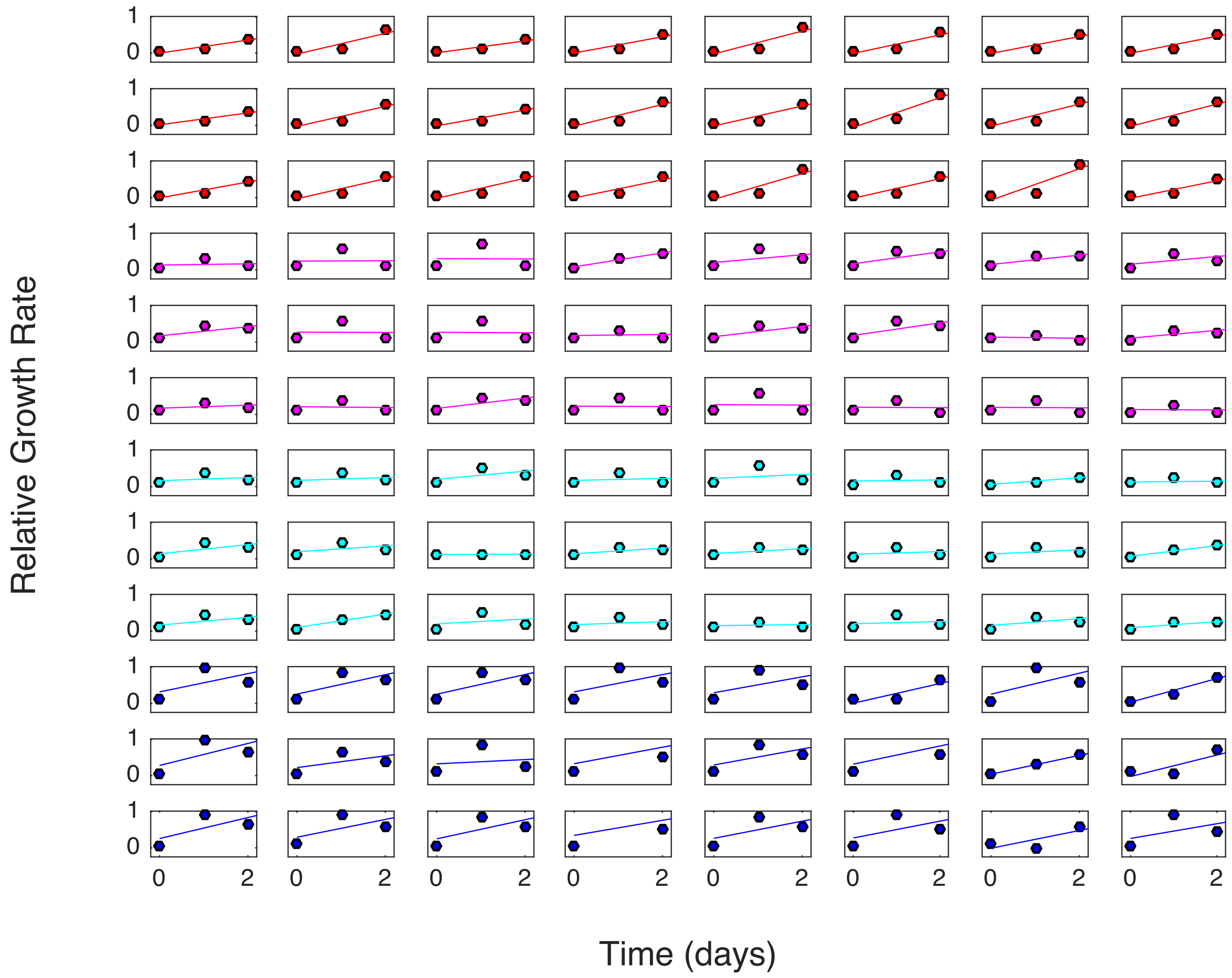


**FIG S2** **Growth rate time series (circles) and linear fits to determine mean adaptation rate (lines) for populations grown in conditions A (top 3 rows, red), B (magenta), C (cyan), and D (last 3 rows, blue) for combinations of streptomycin (STR) and ampicillin (AMP).**

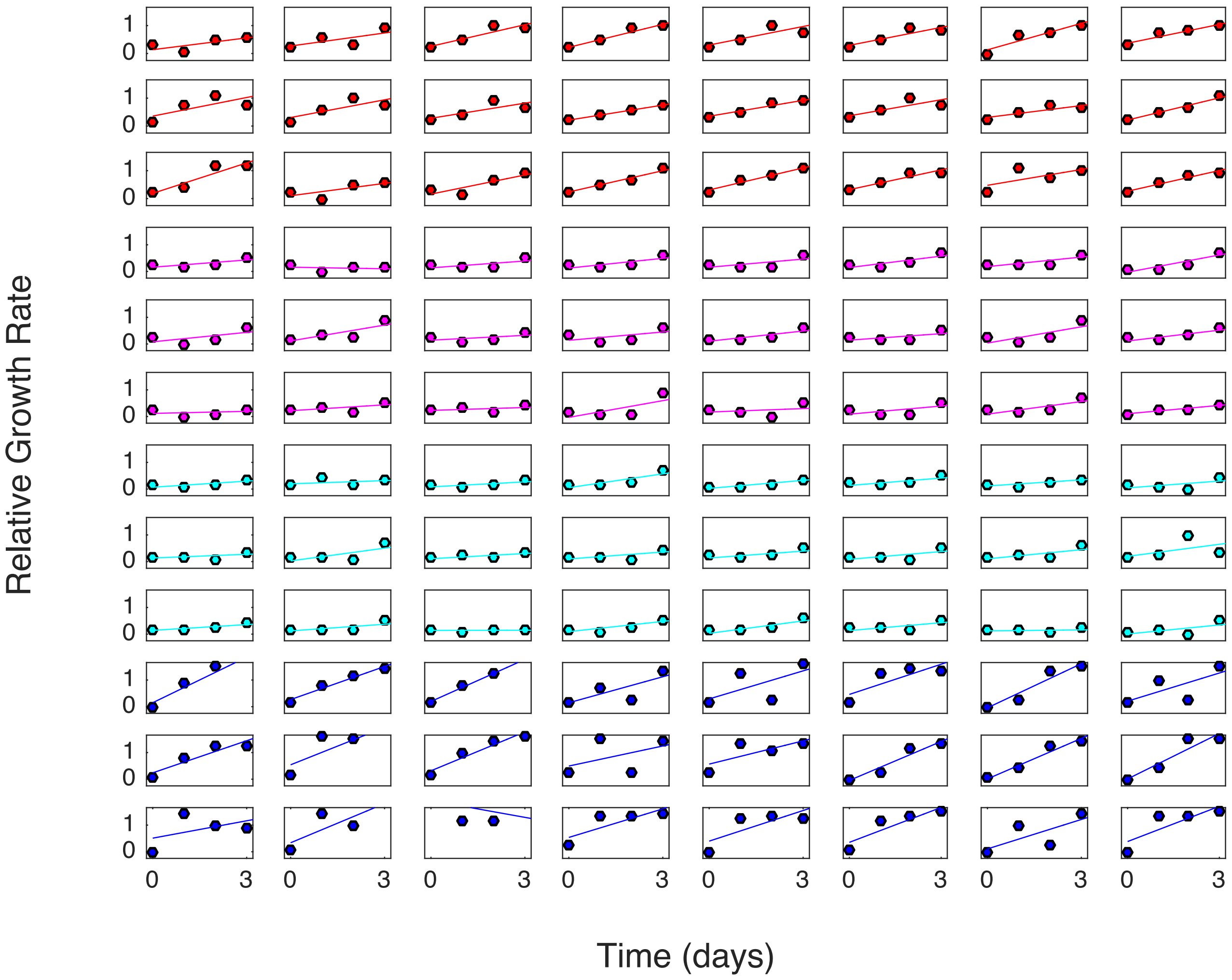


**FIG S3** Growth rate time series (circles) and linear fits to determine mean adaptation rate (lines) for populations grown in conditions A (top 3 rows, red), B (magenta), C (cyan), and D (last 3 rows, blue) for combinations of ceftriaxone (CRO) and ciprofloxacin (CIP).

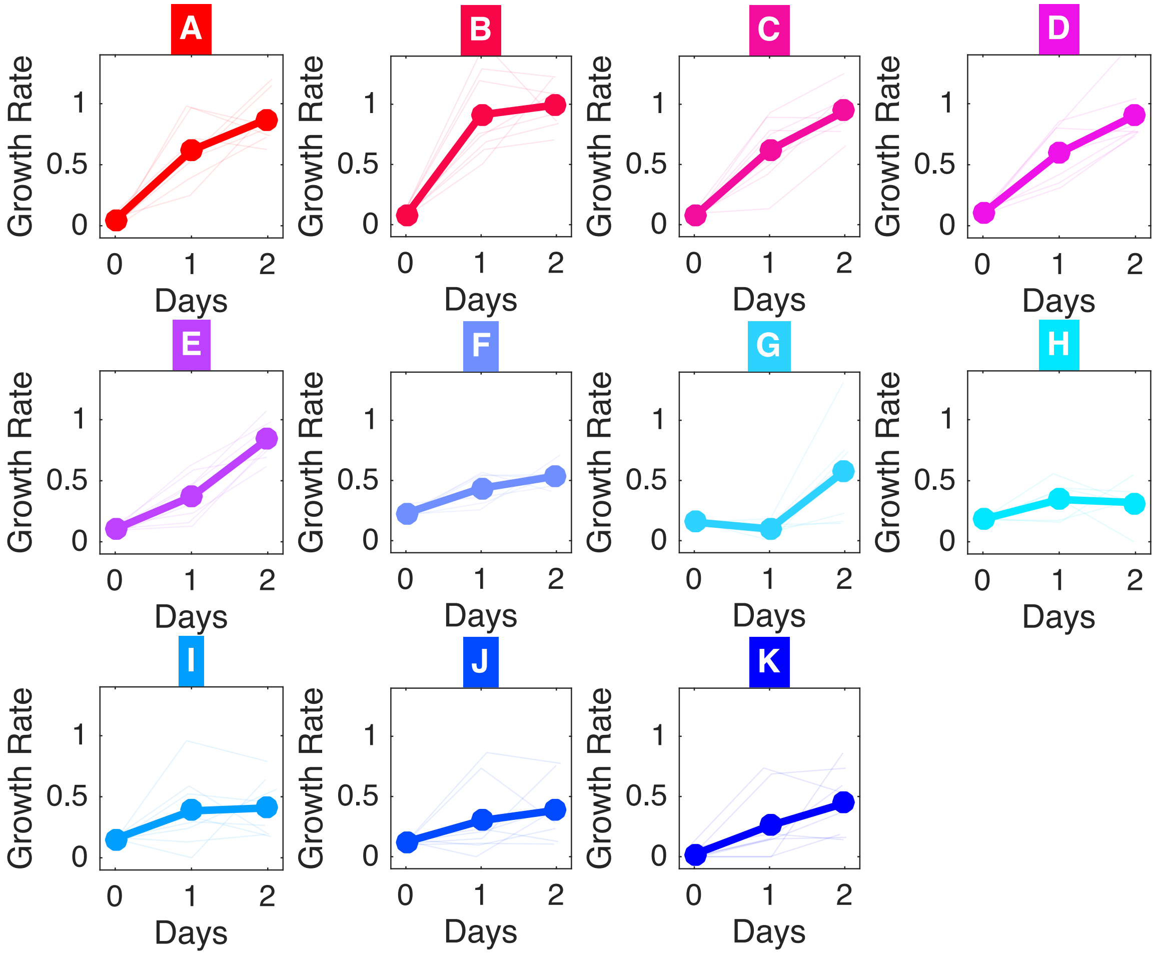


**FIG S4** Growth rate time series for each population (light lines) and the mean across populations for a given condition for combinations of tigecycline (TIG) and ciprofloxacin (CIP).
